# Cryo-EM structure of an atypical proton-coupled peptide transporter: Di- and tripeptide permease C

**DOI:** 10.1101/2022.04.12.487992

**Authors:** Maxime Killer, Giada Finocchio, Haydyn D.T. Mertens, Dmitri I. Svergun, Els Pardon, Jan Steyaert, Christian Löw

**Affiliations:** Centre for Structural Systems Biology (CSSB), Notkestrasse 85, D-22607 Hamburg, Germany; Molecular Biology Laboratory (EMBL), Hamburg Unit c/o Deutsches Elektronen Synchrotron (DESY), Notkestrasse 85, D-22607 Hamburg, Germany; Structural Biology Brussels, Vrije Universiteit Brussel (VUB), Brussels 1050, Belgium; VIB-VUB Center for Structural Biology, VIB, Brussels 1050, Belgium

**Keywords:** peptide transporter, SLC15, proton-dependent oligopeptide transporter (POT), nanobody, macrobody, single particle cryo-EM, DtpC, YjdL

## Abstract

Proton-coupled Oligopeptide Transporters (POTs) of the Major Facilitator Superfamily (MFS) mediate the uptake of short di- and tripeptides in all phyla of life. POTs are thought to constitute the most promiscuous class of MFS transporters, with the potential to transport more than 8400 unique substrates. Over the past two decades, transport assays and biophysical studies have shown that various orthologues and paralogues display differences in substrate selectivity. The *E. coli* genome codes for four different POTs, known as Di- and Tripeptide permeases A-D (DtpA-D). DtpC was shown previously to favor positively charged peptides as substrates. In this study, we describe, how we determined the structure of the 53 kDa DtpC by cryogenic electron microscopy (cryo-EM), and provide structural insights into the ligand specificity of this atypical POT. We collected and analyzed data on the transporter fused to split superfolder GFP (split sfGFP), in complex with a 52 kDa macrobody and with a 13 kDa nanobody. The latter sample was more stable, rigid and a significant fraction dimeric, allowing us to reconstruct a 3D volume of DtpC at a resolution of 2.7 Å. This work provides a molecular explanation for the selectivity of DtpC, and highlights the value of small and rigid fiducial markers such as nanobodies for structure determination of low molecular weight integral membrane proteins lacking soluble domains.

## 1 Introduction

Membranes of cells compartmentalize metabolic processes and present a selective barrier for permeation. To preserve the characteristic intracellular milieu, membrane transporters with specialized functions have evolved to maintain the nutrient homeostasis of cells (Hediger et al., 2013; Zhang et al., 2019). Many of those are energized by an electrochemical proton gradient, providing a powerful driving force for transport and accumulation of nutrients above extracellular concentrations. Proton-dependent oligopeptide transporters (POTs) of the Solute Carrier 15 family (SLC15) are representatives of such secondary active transport systems and occur in all living organisms. They allow an efficient uptake of peptides and amino acids in bulk quantities (Daniel et al., 2006; Thwaites and Anderson, 2007). The best characterized members are the two mammalian PepT1 and PepT2 transporters which are known to play crucial roles in human health, being responsible for the uptake and distribution of nutrients such as di- and tripeptides (Brandsch et al., 2004; Smith et al., 2013; Spanier and Rohm, 2018; Viennois et al., 2018). They also play key roles in human diseases, and impact the pharmacokinetic profiles of orally administered drug molecules (Daniel, 2004; Brandsch, 2009; Ingersoll et al., 2012; Hillgren et al., 2013; Colas et al., 2017; Heinz et al., 2020). SLC15 transporters belong to the Major facilitator superfamily (MFS). MFS transporters share a well-characterized fold, consisting of twelve transmembrane helices organized in two six-helix bundles, expected to function according to the alternate access mechanisms (Jardetzky, 1966) where either side of the transporter is alternately exposed to one side of the membrane. Therefore, substantial conformational changes are required to complete an entire transport cycle with at least three postulated states: (i) inward-open, (ii) occluded, and (iii) outward-open (Yan, 2015; Drew and Boudker, 2016; Quistgaard et al., 2016; Bartels et al., 2021; Drew et al., 2021). POTs have been intensively studied on a structural and biochemical level over the last 30 years. More than 50 entries for this transporter class can be found in the protein data bank, representing ten different bacterial homologues and the mammalian PepT1 and PepT2 transporters, bound to a limited set of substrates and drugs (Newstead et al., 2011; Solcan et al., 2012; Doki et al., 2013; Guettou et al., 2013, 2014; Lyons et al., 2014; Zhao et al., 2014; Quistgaard et al., 2017; Martinez Molledo et al., 2018a, 2018b; Minhas et al., 2018; Nagamura et al., 2019; Ural-Blimke et al., 2019; Killer et al., 2021; Parker et al., 2021; Stauffer et al., 2022). Although bacterial and eukaryotic POTs share an overall conserved binding site, individual amino acids changes in or in close vicinity of the binding site are likely responsible for observed differences in affinities and selectivity for particular peptides and drugs among the studied POT homologues. Here, structural biology studies are particularly crucial to understand substrate promiscuity and drug coordination on a molecular level. While bacterial POT structures, determined by mainly X-ray crystallography, represent exclusively the inward-open or inward-open-partially occluded state, the mammalian PepT1 and PepT2 transporters were recently captured in various conformations by single particle cryo-EM, advancing the mechanistic understanding of the entire transport cycle (Killer et al., 2021; Parker et al., 2021). Despite their small size of typically only 50 kDa for an individual transporter unit, these systems become more and more accessible for single-particle cryo-EM approaches. Indeed, in 2021, more MFS transporter structures were determined by single-particle Cryo-EM (17 pdb entries; resolution range 3.0 – 4.2 Å) than X-ray crystallography (14 pdb entries; resolution range 1.8 – 3.6 Å).

Although known POT structures show a high level of similarity, various works have indicated that homologues can differ in their range of transported substrate and drug molecules (Lyons et al., 2014; Prabhala et al., 2014; Boggavarapu et al., 2015; Sharma et al., 2016; Martinez Molledo et al., 2018a). The *E. coli* genome codes for four different POTs named Di-and Tripeptide permease A-D (DtpA-D), also known as YdgR (=DtpA), YhiP (=DtpB), YjdL (=DtpC) and YbgH (=DtpD). They cluster in pairs, DtpA and B (sequence identity 51%), and DtpC and D (sequence identity 56%) with around 25% identity between them. DtpA and B exhibit a prototypical substrate preference similar to the human PepT1 transporter (Chen et al., 2000; Harder et al., 2008; Foley et al., 2010; Prabhala et al., 2017, 2018), while DtpC and D have been classified as atypical POTs, because DtpC prefers di-peptides in particular those with a lysine residue in the second position. Although DtpC has been well characterized in terms of function over the last years (Ernst et al., 2009; Jensen et al., 2012c, 2012a, 2012b, 2014; Prabhala et al., 2014; Aduri et al., 2015), it has resisted structure determination by X-ray crystallography so far (Gabrielsen et al., 2011).

Here we describe the structure determination of the bacterial POT transporter DtpC by single particle cryo-EM. Considering that the transporter displays no characteristic cytoplasmic or periplasmic features which are helpful to drive the particle alignment, we applied different strategies previously described in the literature to increase the overall size of the transporter to overcome these limitations. We i) fused the transporter to split-sfGFP (Liu et al., 2020, 2022), ii) raised different nanobodies against DtpC (Pardon et al., 2014) and iii) extended the nanobody to a macrobody (Brunner et al., 2020; Botte et al., 2021). The various samples were subsequently imaged by cryo-EM and analysed. DtpC in complex with the conformation specific nanobody 26 turned out to be more rigid and a significant fraction of the sample dimeric, allowing us to reconstruct DtpC to 2.7 Å resolution. The DtpC structure now provides molecular insights into how selectivity within this transporter family is achieved.

## 2 Results and Discussion

### Different fiducial marker strategies for structure determination

Since MFS transporters typically lack additional domains outside their transport unit, which is a major impediment for accurate particle alignment in single particle cryo-EM approaches, we assessed three fiducial marker strategies introducing additional density outside of detergent micelles containing DtpC, by analyzing the quality of 2D class averages (Fig 1). To obtain conformation specific binders against DtpC, we first immunized llamas with recombinant DtpC and selected nanobodies (Nbs) following standard procedures (Pardon et al., 2014). Three out of five selected binders (Nb17, Nb26, and Nb38) co-eluted with DtpC on gel filtration and increased the melting temperature of the respective DtpC-Nb complex by 20 °C, 16 °C, and 12 °C. (Fig 2 A,B). DtpC in complex with Nb17 and Nb26 yielded crystals in various conditions, but despite extensive optimization efforts, the crystals of the DtpC-Nb26 complex did not diffract X-rays better than 5 Å resolution. In a second step, we decided to increase the size of Nb26, which formed a tight complex with DtpC, by fusing one copy of the maltose binding protein (MBP) to its C-terminus as described previously (Botte et al., 2021). This resulted in a 52 kDa macrobody, and we expected it to bind to the periplasmic side of the transporter as seen in other MFS transporter-Nb complexes (Fig 1). In a third approach, we fused the two self-assembling parts of split-sfGFP; with β1-6 on the N-terminus of DtpC, and β7-11 on the C-terminus. We named this construct split sfGFP-DtpC_FL_. In order to minimize the mobility between the membrane protein and the split sfGFP fiducial, we also generated two additional constructs where the last five (split sfGFP-DtpC_1-475_), or ten residues (split sfGFP-DtpC_1-470_) of the transporter were deleted. We then assessed proper folding and complementation by monitoring the fluorescence of the chromophore on an HPLC system (Fig 2C). All constructs eluted at similar retention times and the fluorescence was highest in the non-truncated construct (split sfGFP-DtpC_FL_) and lowest in the most truncated version (split sfGFP-DtpC_1-470_). In order to extend this observation to other MFS transporters, we repeated this experiment with the human POT homologue PepT1, and noticed a similar trend upon shortening of the termini. Yet, since the decrease of fluorescence was only minor in split sfGFP-DtpC_1-475_ in comparison to split sfGFP-DtpC_FL_, we proceeded to imaging with the shorter construct in the presence of Nb26.

**Figure 1:**
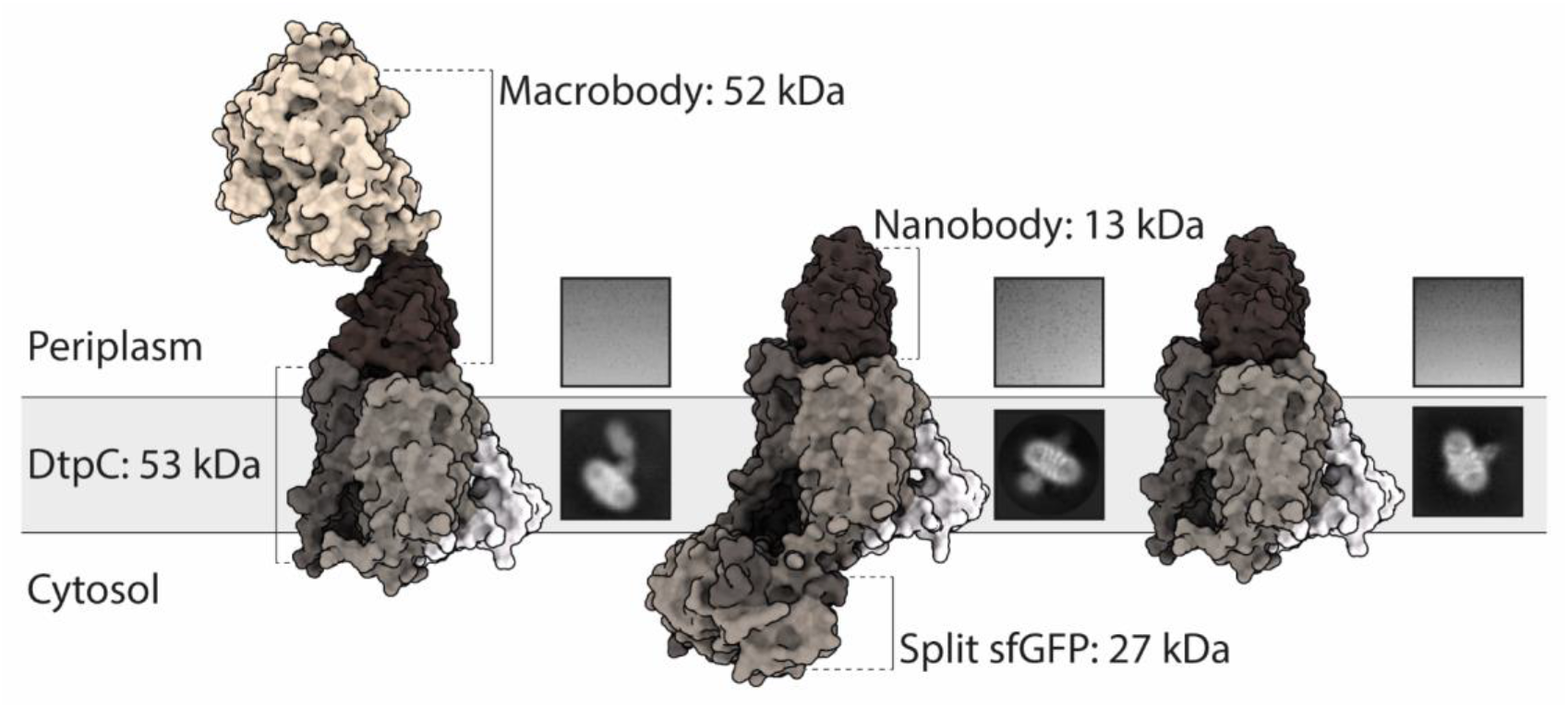
Utilization of different fiducial markers to improve particle alignment and 2D averaging from cryo-EM images. From left to right: DtpC-Mb26, split-sfGFP-DtpC_1-475_-Nb26, and DtpC-Nb26 were purified, vitrified on grids and imaged. Single particles were identified, clustered and averaged. The best average from each sample is shown under a representative raw micrograph.

**Figure 2:**
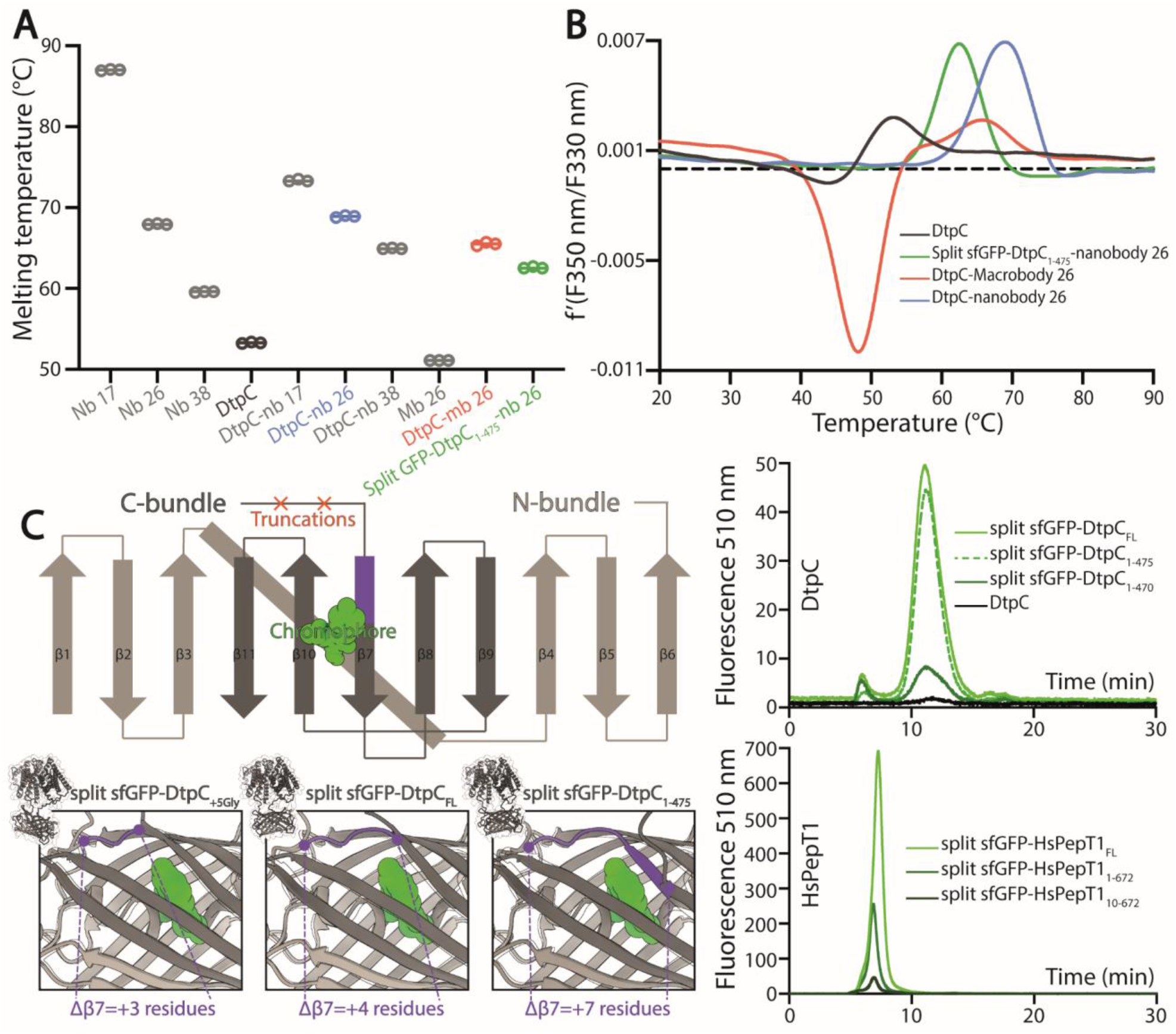
Characterization of the different fiducial markers. (A) The melting temperature of each fiducial and DtpC-fiducial complex was measured by nano-differential scanning fluorimetry (DSF) in triplicate measurements as shown as open circles. The average of the three values is marked by a line. (B) The first derivative of the summarized data in (A) is shown for DtpC and the three imaged samples together with the respective fiducial. (C) A schematic representation of the split sfGFP-DtpC architecture is shown on the top left panel. Below, structure predictions were generated for split-sfGFP-DtpC_+5Gly_, split-sfGFP-DtpC_FL_, sfGFP-DtpC_1-475_, and overlaid with sfGFP (PDB accession number 2B3P). The dark-violet coloring corresponds to the fraction of ß7 which is properly folded in sfGFP while unfolded in the restrained chimeric construct. The right panel shows HPLC chromatogram profiles monitoring the fluorescence of the chromophore of split sfGFP in the context of the indicated constructs, using 480 nm as excitation wavelength and recording at 510 nm the emitted light.

The particle density and distribution in the vitrified solution was similar in the three imaged samples. However, DtpC-Nb26 produced the best 2D class averages considering the sharpness of secondary structure elements inside the micelle, as judged by visual inspection (Fig 1, Fig S1). The Nb26-MBP (Mb26) fiducial was clearly visible in 2D class averages, but it adopted various positions in relation to the transporter, therefore making accurate alignment of the particles more difficult than in its shorter but more rigid and stable nanobody version (Fig 1, Fig 2 A,B, Fig S1). The split sfGFP-DtpC_(1-475)_-Nb26 sample allowed clear visualization of the transmembrane helices after clustering a small subset of particles, but the majority of particles clustered in classes with blurry density for the split sfGFP fiducial, or with the two complementary parts β1-6 and β7-11 not assembled (Fig S1). Alphafold2 predictions on the imaged construct, as well as on the full length construct later suggested a destabilization of the beta-barrel upon increasing termini restrains, resulting in partial unfolding of β7 and exposure of the chromophore to solvent quenching. Interestingly, this effect could partially be reverted by adding a linker of five glycine residues between the C-bundle and β7 based on *in silico* data. We conclude that termini restraining using the split-sfGFP approach is a promising fiducial strategy for structural studies of MFS transporters, in addition to the previous demonstrated showcases on small membrane proteins (2, 4 and 6TMs) (Liu et al., 2020, 2022). However, the amount of restraining in larger membrane proteins such as MFS transporters where both termini are placed far from each other need to be optimized experimentally or *in silico*, to produce a stable and rigid fiducial; two crucial aspects for high resolution structure determination of MFS transporters by single particle cryo-EM.

As we obtained the best 2D class averages for DtpC with the fiducial marker Nb26, we proceeded to a large data collection (Table 1) and could cluster a subset of dimers within this data set. The presence of different oligomeric species was already expected based on the peak shape of the gel filtration run. The large mass of the dimer, and the stable and rigid signal of the Nb26 fiducial, allowed us to reconstruct the DtpC-Nb26 dimer to 3.0 Å resolution and model this assembly (Fig 3, 4, Fig S2). The quaternary structure consists of a non-symmetrical inverted dimer mediated by interactions through a large hydrophobic interface between the HA-HB helices of DtpC (Fig S2). Although other inverted dimers were reported in homologous POT structures (Quistgaard et al., 2017), the source of such arrangements is likely to be artificial. We also investigated the oligomer heterogeneity in solution with small angle X-ray scattering and obtained a good fit at low angles (corresponding to the overall shape of particles in solution) for the cryo-EM volume of the dimer (Fig S3). The fit to a monomeric cryo-EM volume was poor, indicating that in detergent solution a significant fraction of DtpC-Nb26 is dimeric. As for the interaction between the membrane protein and the fiducial marker, the CDR3 loop of Nb26 accounts for the strongest interactions with the periplasmic surface of the transporter with two salt bridges, while CDR1 and CDR2 contribute *via* hydrogen bounding (Fig 5). 3D variability analysis (Punjani and Fleet, 2021) revealed a small degree of flexibility between the two DtpC-Nb26 copies. Therefore, we performed a local refinement, focused on one copy of the membrane protein, which extended the resolution of the reconstruction to 2.7 Å and improved the accuracy of the atomic model for subsequent structural analysis (Fig 4).

**Table 1:** Data collection and refinement statistics of the deposited DtpC structure

|  |  |
| --- | --- |
| **Protein reconstructed** | Di- and tripeptide permease C (DtpC) |
| **PDB accession code** | 7ZC2 |
| **EMDB accession code** | EMD-14618 |
| **Data acquisition** |  |
| Microscope/Detector | Titan Krios/Gatan K3 |
| Imaging software | EPU |
| Magnification | 105,000 |
| Voltage (kV) | 300 |
| Electron exposure (e-/Å<sup>2</sup>) | 75 |
| Dose rate (e-/pix/s) | 19.5 |
| Frame exposure (e-/Å<sup>2</sup>) | 1.5 |
| Defocus range (μm) | -0.9 to -1.8 |
| Physical pixel size (Å) | 0.85 |
| Micrographs | 24,333 |
| **Reconstruction** |  |
| Picked coordinates (cryolo) | 6,464,070 |
| Particles in 3D classification (RELION) | 6,365,235 |
| Particles in final refinement (CryoSPARC) | 878,428 |
| Symmetry imposed | C1 |
| Map sharpening method | Phenix Resolve_cryo_em |
| Map resolution, FSC<sub>half maps</sub>; 0.143 masked/unmasked (Å) | 2.72/3.43 |
| **Refinement** |  |
| Initial model used for refinement | AlphaFold2 model, relaxed with Amber |
| Model resolution (Å) |  |
| FSC 0.143, masked/unmasked | 2.64/5.43 |
| Model composition |  |
| Non-hydrogen atoms | 7334 |
| Protein residues | 471 |
| ADP B factor (Å<sup>2</sup>) mean | 12.73 |
| R.m.s deviations |  |
| Bond lengths (Å) (#>4σ) | 0.003 (0) |
| Bond angles (°) (#>4σ) | 0.616 (0) |
| Validation |  |
| MolProbity score | 1.44 |
| Clashscore | 8.04 |
| Rotamer outliers (%) | 0.00 |
| Ramachandran plot |  |
| Favored (%) | 98.29 |
| Allowed (%) | 1.71 |
| Outliers (%) | 0.00 |

**Figure 3:**
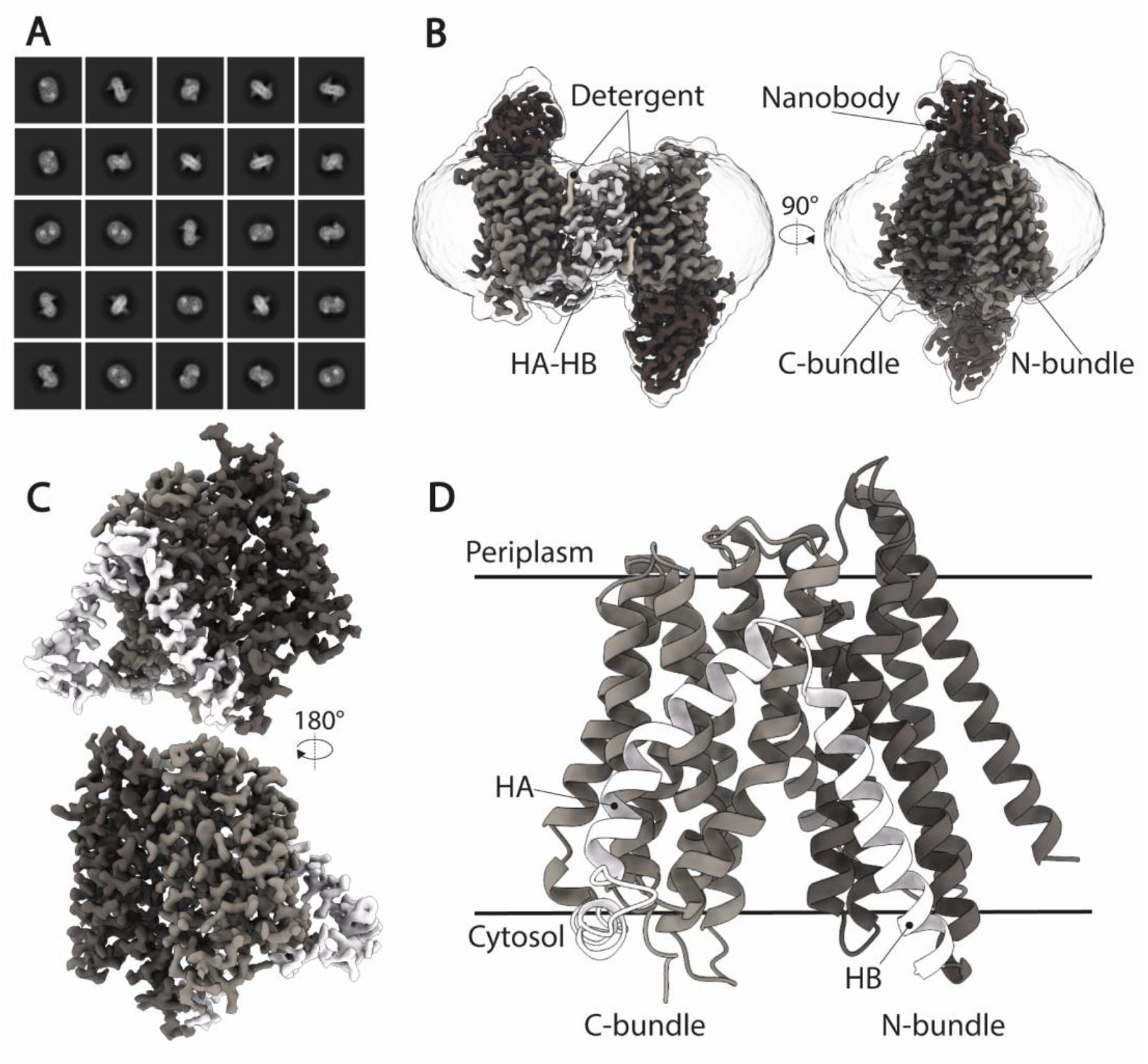
Cryo-EM structure of DtpC-Nb26. (A) Representative 2D class averages of the dimeric population. (B) 3D reconstruction of the DtpC-Nb26 inverted dimer used for local focused refinement on one copy of the transporter, shown in (C). (D) Atomic model of DtpC displayed as ribbon diagram. The different structural elements are labelled.

**Figure 4:**
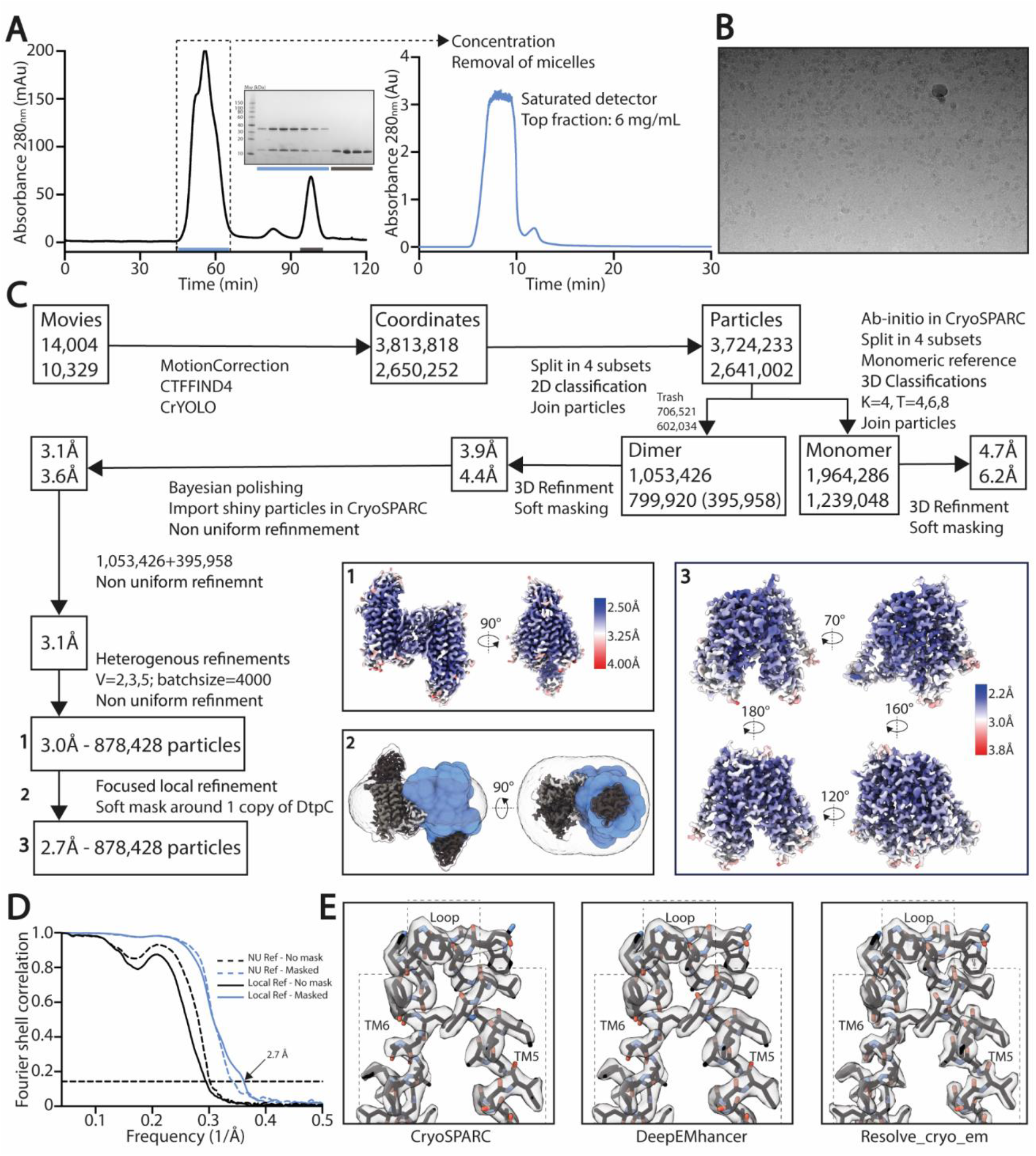
High resolution structure determination of DtpC-Nb26. (A) Gel filtration was performed on a preparative column (left) before concentrating the sample to 60 mg/mL and rerunning it on an analytic column on an HPLC system (right), in order to obtain a highly concentrated sample, free of empty detergent micelles. Peak shape already indicates a mixture of different oligomeric species. (B) Representative raw micrograph of the acquired dataset. The applied defocus is −1.5 µm. (C) Summary of the image analysis. The angular assignments from the dimeric reconstruction were used as prior to perform a local focused refinement with reduced angular and translational searches on the masked region illustrated in blue. (D) The Fourier transforms over different shells on frequency space, of two independent volumes (half maps) were compared (FSC) and plotted as a function of spatial frequency, to estimate the overall resolution using the 0.143 cutoff threshold. (E) The two half maps were used as inputs to assess various post-processing strategies.

**Figure 5:**
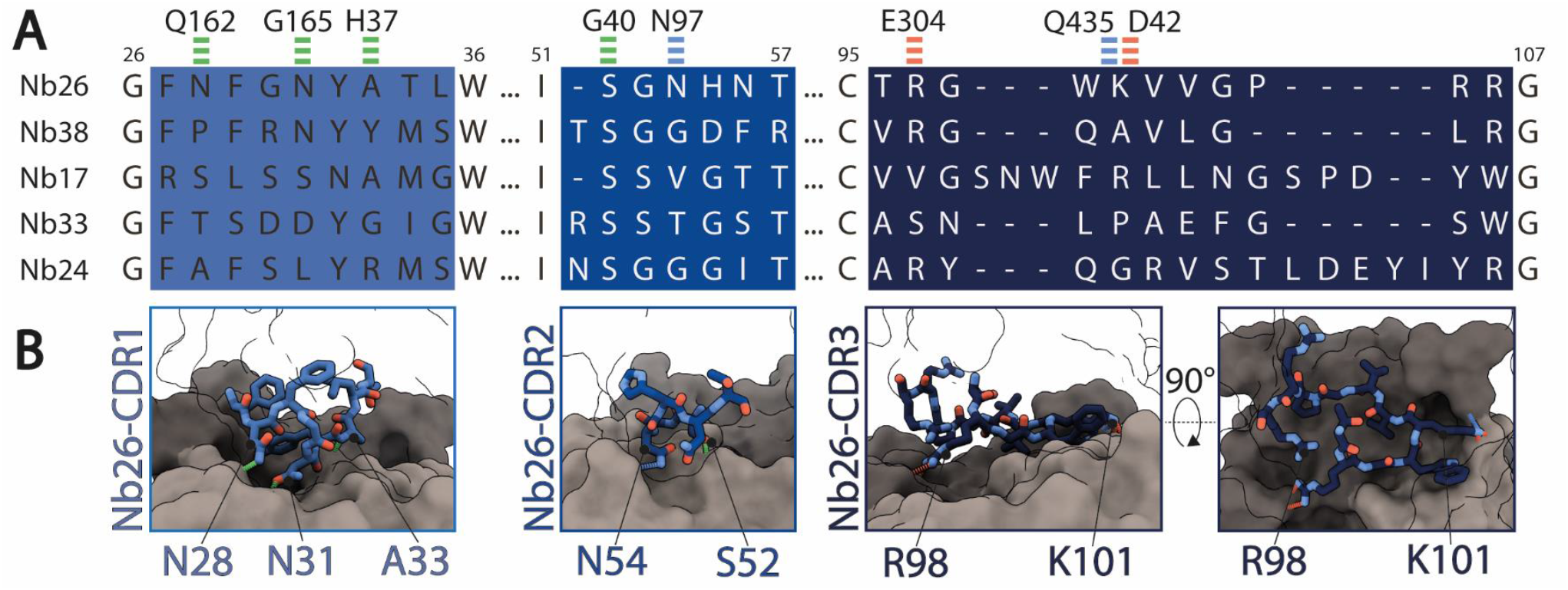
Interactions between Nb26 and DtpC. (A) The sequences of the five nanobodies representing five different families, obtained after selection, with their complementary determining regions (CDR) are shown. Interactions of Nb26 with DtpC are highlighted as green (hydrogen bounds involving the protein backbone), blue (hydrogen bounds involving side chains) and red dashes (salt bridges). (B) These interactions are further displayed in 3D. CDR regions are depicted as sticks on the surface of DtpC where the N-terminal bundle is colored in grey, and the C-terminal bundle in dark grey.

### Structural basis for ligand selectivity in DtpC

The DtpC structure revealed the expected and well-known MFS transporter fold, with twelve transmembrane helices (TMs) organized in two helical bundles and additional two TMs specific for the POT family (known as HA and HB domains). The peptide binding site of DtpC is exposed to the cytoplasmic side (Fig 3, 4). Almost all bacterial POT structures described so far were determined by X-ray methods in a similar inward facing (IF) conformation. The extent to which the central cavity is open to the cytosol is regulated by a mechanism of occlusion mediated by TM4, TM5, TM10, and TM11, as supported by structures in IF occluded, partially occluded, and open states. In the case of the here described DtpC structure, the IF state is open (Fig 3).

Molecules from the periplasmic side, on the contrary, cannot enter the central cavity. Tight closure of both bundles above the binding site is mediated by a salt bridge between D43 (TM2, N-bundle) and R294 (TM7, C-bundle) and hydrogen bounds between H37 (TM1, N-bundle) and D293 (TM7) as well as R28 (TM1) and N421 (TM11, C-bundle) (Fig 6A). We also analyzed previously determined POT structures with clearly resolved side chain densities, to understand how the IF state is generally maintained in this transporter family. Except for human PepT2 and the POT transporter from *Shewanella oneidensis* (PepT_So_), where the inter-bundle periplasmic salt bridge is formed between TM5 and TM7, the IF state is in all other analyzed structures stabilized by a salt bridge on the tip of TM2 and TM7 (Fig 6B). Additional hydrogen bonding networks as described in other studies, can occur, but vary greatly among different homologues. This analysis highlights that the alternate access mechanism in canonical and in so called ‘atypical’ POTs share similarities such as electrostatic clamping by formation and disruption of salt bridges. The differences in hydrogen bonding patterns however, could account for the various turnover rates seen among POT homologues.

**Figure 6:**
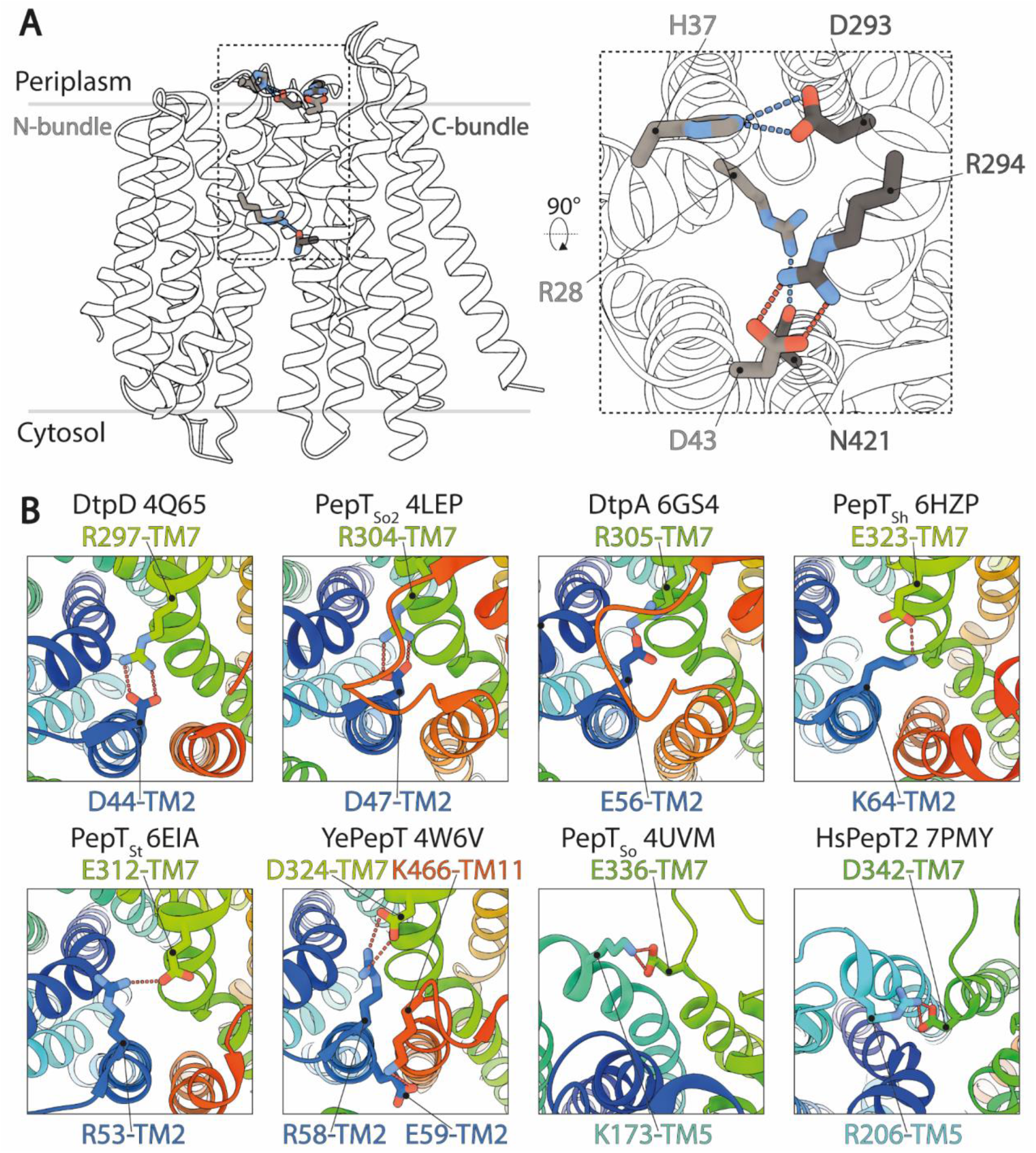
Structural basis for the stabilization of the inward facing state in DtpC and other POT homologues. (A) The salt bridge and hydrogen bounds favoring closure of the two bundles on the periplasmic side of DtpC are respectively shown as red and blue dashes. (B) The structures of homologous POTs from *Escherichia coli* (DtpD, DtpA), *Shewanella oneidensis* (PepT_So2_, PepT_So_) *Staphylococcus hominis* (PepT_Sh_), *Streptococcus thermophilus* (PepT_St_), *Yersinia enterocolitica* (YePepT) and *Homo sapiens* (HsPepT2), were all previously captured in the IF state. Here they were analyzed to identify the strongest interaction stabilizing their common conformation. The structures are colored from blue to red, from their N-to C-termini, and the respective PDB accession numbers are indicated. Conserved salt bridges are labelled and highlighted by red dashed lines.

Canonical POTs are characterized by i) the presence of the E_1_XXE_2_R motif on TM1 involved in proton coupling and ligand binding, and ii) the ability to accommodate dipeptides, tripeptides, and peptidomimetics, which relies on a set of conserved residues located in the central binding cavity. In DtpC, the E_1_XXE_2_R motif, has evolved to Q_1_XXE_2_Y (where Q_1_=N17, E_2_=E20, Y=Y21). In all high resolution X-ray structures of canonical POTs, R is in salt-bridge distance to E_2_ and the C-terminus of substrate peptides. Mutation of either E_1_ or E_2_ in the conventional E_1_XXE_2_R motif to glutamine residues abolishes uptake (Aduri et al., 2015). A reverse mutation in DtpC, from Q_1_XXE_2_Y to E_1_XXE_2_Y or to E_1_XXQ_2_Y preserves high transport rates, while a Q_1_XXQ_2_Y motif significantly decreases it (Aduri et al., 2015). In addition, based on previous molecular dynamics experiments, a salt bridge switching mechanism from R-E_2_ to R-E_1_, upon protonation of E_2_ in the E_1_XXE_2_R motif, was proposed (Aduri et al., 2015). This biochemical and *in silico* data strongly support a dual role of the E_1_XXE_2_R motif for both proton and peptide transport, where R can form a salt bridge interaction with the C-terminus of peptides or with E_1_ when E_2_ is protonated, and where the deprotonation event of the latter is required to disrupt the R-peptide interaction.

In DtpC, we now observe that the side chain pocket has a different architecture and characteristic in comparison with the one of canonical POTs. It displays an overall more acidic groove caused by the presence of the aspartate residue 392. Canonical POTs have a conserved serine residue instead, yielding a slightly changed hydrophobicity pattern in the binding site (Fig 7 A-D). A structural overlay of DtpC with a canonical POT structure bound to the dipeptide Ala-Phe allows us to position the peptide in the binding site. By replacing the phenyl group with a lysine side chain (generating the known DtpC dipeptide substrate Ala-Lys instead of Ala-Phe), we postulate a putative salt bridge between the carboxyl group of D392 and the ε-amino group of the lysine side chain. This observation, together with previous biochemical work (Jensen et al., 2012b; Aduri et al., 2015) allows us to hypothesize that the selectivity of DtpC for dipeptides with C-terminal lysine or arginine residues is caused by swapping a salt bridge between the recurrent carboxyl group of the peptide terminus and the transporter (R21Y mutation), to a side chain specific salt bridge with D392. Since the R-peptide interaction is lost in DtpC, there is no requirement for E1 to destabilize R-peptide for release, which would explain the presence of a Q_1_XXE_2_Y motif instead of E_1_XXE_2_R.

**Figure 7:**
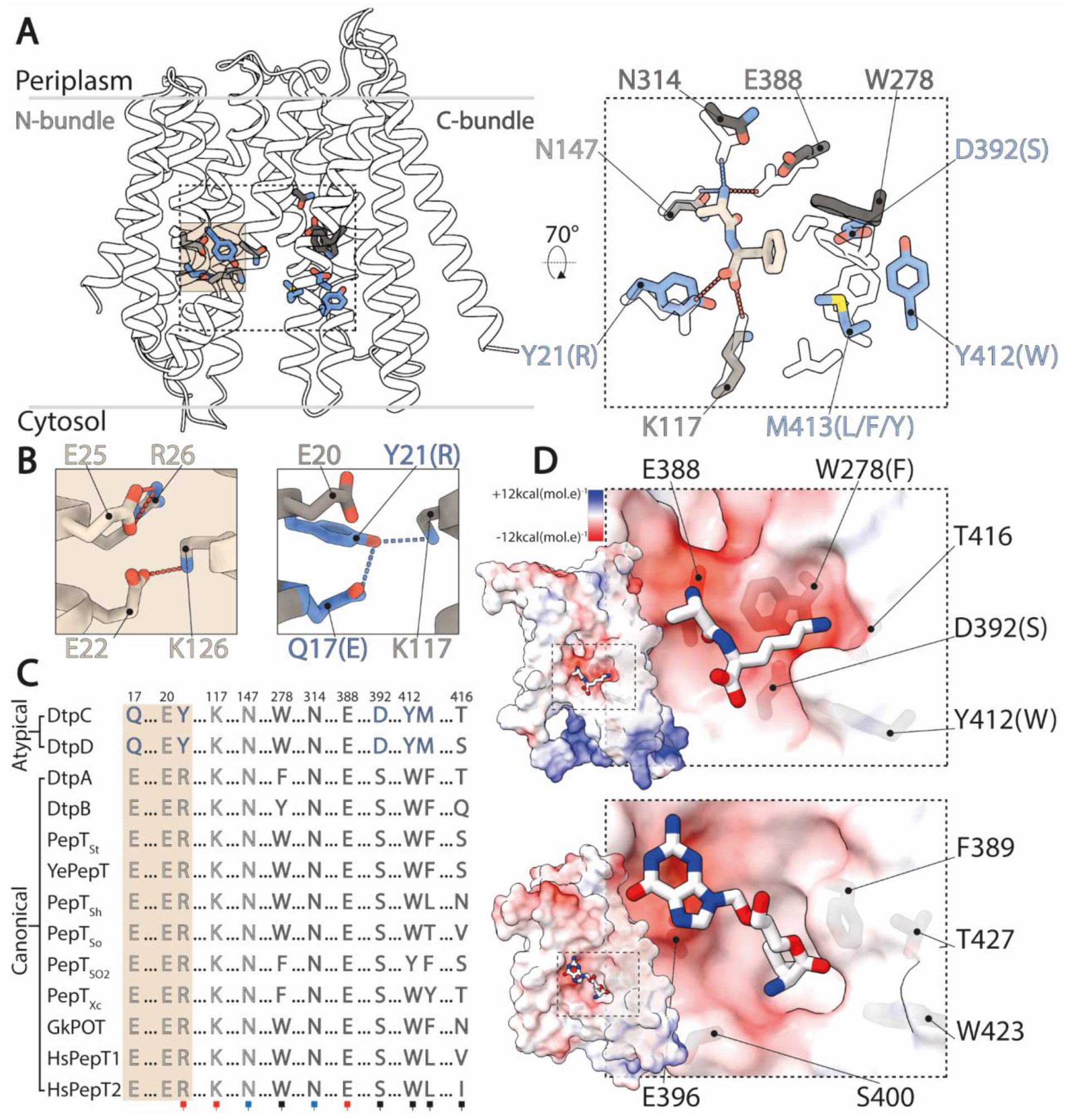
Structural basis for ligand selectivity in DtpC and atypical POTs. (A) Key residues involved in substrate binding are colored and shown as sticks. In the close up view, an overlay of HsPepT2 (transparent residues) bound to the dipeptide Ala-Phe (beige) with DtpC is shown. Residues colored in grey are conserved while blue residues are exclusive to atypical POTs. (B) The arrangement of the E1XXE2R motif from PepT_St_ is shown on the left panel, and the atypical Q1XXE2Y on the right. (C) The sequences of 13 POTs were aligned and residues involved in proton coupling and substrate binding are shown. The red squares mark residues strongly interacting with the charged termini of substrates peptides via salt bridges. The blue squares indicate two conserved asparagine residues stabilizing peptides through hydrogen bounds. The black squares point to residues constituting the side chain pocket of POTs, tuning ligand promiscuity or selectivity. (D) Surface representation colored by electrostatic potential, of the C-bundles of DtpC (top panel) and DtpA (bottom panel). A pose of the preferred substrate of DtpC, Ala-Lys, is proposed (top) and the co-crystallized valganciclovir drug hijacking canonical POTs is shown in DtpA (bottom). PDB accession codes of previously published work: HsPepT2 bound to Ala-Phe: 7PMY; PepT_St_: 5OXO; DtpA bound to valganciclovir: 6GS4.

In summary, our work provides new insights into promiscuous *versus* selective substrate recognition in POTs and constitutes a step forward towards completing the family of *E. coli* POTs structures. Lastly, it displays some of the challenges related to high resolution cryo-EM structure determination of MFS transporters devoid of soluble domains, and manifests once again, the benefit of fiducial markers in overcoming those.

## 3 Material and Methods

### Expression and purification of membrane protein constructs: DtpC; split sfGFP-DtpC (full length split sfGFP-DtpC_FL_, and truncated constructs split sfGFP-DtpC_1-475_ and split sfGFP-DtpC_1-470_); split sfGFP-*Hs*PepT1 (full length split sfGFP-*Hs*PepT1_FL_, and truncated constructs split sfGFP-*Hs*PepT1_1-672_ and split sfGFP-*Hs*PepT1_10-672_)

The full-length cDNA of DtpC wild type (WT) was amplified from the *Escherichia coli* genome, and cloned into a pNIC-CTHF vector by ligation-independent cloning (LIC). This vector contains a C-terminal His-Tag and a Tobacco Etch virus (TEV) cleavage site and a kanamycin resistance gene as selectable marker. The first 6 N-terminal beta strands of sfGFP were fused to the N-terminus of DtpC, and the beta strands 7 to 11 fused to the C-terminus. We named this construct split sfGFP-DtpC_FL_. Two additional constructs were cloned with truncations of 5 (split sfGFP-DtpC_1-475_), and 10 residues (split sfGFP-DtpC_1-470_), on the C-terminal side of DtpC.

*Hs*PepT1 was previously cloned into a pXLG vector containing an expression cassette composed of an N-terminal Twin-Streptavidin tag followed by the HRV-3C protease recognition sequence (Killer et al., 2021). Similarly, as for DtpC, the two self-assembling parts of split-sfGFP were first inserted into the N- and C-termini of the full-length version of *Hs*PepT1, and on two other versions with i) a C-terminal truncation of 36 residues (split sfGFP-HsPepT11-672), and ii) a C-terminal truncation of 36 residues and a N-terminal truncation of 10 residues (split sfGFP-HsPepT110-672) were cloned.

Recombinant DtpC, and the three split sfGFP-DtpC constructs were expressed in *E. coli* C41(DE3) cells grown in terrific broth (TB) media supplemented with 30 μg/ml kanamycinaccording to established procedures (Löw et al., 2012, 2013). Cultures were grown at 37°C and protein expression was induced with 0.2 mM IPTG at an OD_600 nm_ of 0.6 - 0.8. After induction, culture growth continued at 18°C for 16-18 hours. Cells were harvested by centrifugation (10,000 g, 15 minutes, 4°C), and the pellet was stored at −20°C until further use. Cell pellets were resuspended in lysis buffer (20 mM NaPi at pH 7.5, 300 mM NaCl, 5% (v/ v) glycerol, 15 mM imidazole, with 3 ml of lysis buffer per gram of wet weight pellet), supplemented with lysozyme, DNase and 0.5 mM tris(2-carboxyethyl)phosphine (TCEP). The cells were lysed by three cycles using an Avestin Emulsiflex homogenizer at 10.000- 15.000 psi. Recovered material was centrifuged to remove non-lysed cells (10,000 g, 15 minutes, 4°C) and the supernatant was subjected to ultracentrifugation to separate the membrane fraction (100,000g, 1 hour, 4°C using an Optima XE-90, Beckman Coulter centrifuge). Membranes were resuspended in lysis buffer supplemented with cOmplete EDTA-free protease inhibitors (Roche), and solubilized by adding 1% n-dodecyl-β-D-maltoside (DDM) detergent (Anatrace). The sample was centrifuged for 50 min at 90,000g, and the supernatant was applied to Ni-NTA beads for immobilized-metal affinity chromatography (IMAC) on a gravity column. The beads were pre-equilibrated in lysis buffer and incubated with the solubilized membrane proteins for one hour at 4°C on a rotating wheel. Loaded beads were washed with buffer with increasing imidazole concentrations (20 mM NaPi at pH 7.5, 300 mM NaCl, 5% glycerol, 15-30 mM imidazole, 0.5 mM TCEP, 0.03% DDM). The proteins were eluted from the column with a buffer containing high imidazole concentration (20 mM NaPi at pH 7.5, 150 mM NaCl, 5% glycerol, 250 mM imidazole, 0.5mM TCEP, 0.03% DDM) and combined with 1 mg of TEV protease to perform the His-tag cleavage during dialysis overnight at 4°C. The dialysis buffer contained 20 mM HEPES at pH 7.5, 150 mM NaCl, 5% glycerol, 0.5 mM TCEP, 0.03% DDM. The cleaved protein was recovered by negative IMAC, concentrated to 4 ml using a 50 kDa concentrator (Corning® Spin-X® UF concentrators) and run on an ÄKTA Pure system (GE Healthcare Life Sciences), using a HiLoad 16/ 600 Superdex 200 column for DtpC, and a Superdex 200 Increase 10/300 column for the split sfGFP-DtpC constructs. Fractions containing the protein were pooled, concentrated, flash frozen and stored at −80°C until further use.

For the split sfGFP-*Hs*PepT1 constructs, expression was done in mammalian cells as described previously (Pieprzyk et al., 2018; Killer et al., 2021). Briefly, HEK293F cells were collected 48 hours after transient transfection, and stored at −80°C until further use. Frozen cell pellets were resuspended in 300 mM NaCl, 20 mM NaPi (pH 7.5), 0.5 mM TCEP, and 5% glycerol, supplemented with cOmplete EDTA-free protease inhibitors, and were disrupted using an Avestin Emulsiflex homogenizer at 10,000-15,000 psi. The lysate was centrifuged for 10 min at 10,000g, 4°C, and the supernatant was centrifuged for 90 min at 100,000g, 4°C. The pellet containing the membrane fraction was solubilized in 1% N-dodecyl-β-D-maltopyranoside (DDM) and 0.1% cholesteryl hemisuccinate (CHS; Tris Salt, Anatrace) for 1 hour at 4°C. The sample was centrifuged for 50 min at 90,000g, and the supernatant was applied to Strep-TactinXT beads (IBA). After 20 min of incubation on a rotating wheel, the suspension was transferred to a gravity column. Following two wash steps with 300 mM NaCl, 20 mM HEPES (pH 7.5), 0.03% DDM, and 0.003% CHS, split sfGFP-*Hs*PepT1comnstructs were eluted with 0.03% DDM, 0.003% CHS, 150 mM NaCl, 20 mM HEPES (pH 7.5), and 10 mM desthiobiotin (Sigma-Aldrich).

### Selection, expression and purification of nanobodies against DtpC

To generate DtpC specific nanobodies, two non-inbred llamas were injected six times at weekly intervals with a mixture of 94 different proteins including DtpC (50 µg of each antigen weekly). After six weeks of immunization, two separate phage display libraries were constructed, one from each animal, in the pMESy2 vector, which is a derivative of pMESy4 that contain a C-terminal EPEA-tag for affinity purification. After pooling both libraries, nanobodies were selected against individual antigens in two rounds of parallel panning in 96-well plates containing one immobilized antigen in each well. After two selection rounds on DtpC, 60 clones were picked for sequence analysis, 13 clones encoded antigen-specific nanobodies as tested in ELISA, grouping them in 5 different sequence families. A nanobody family is defined as a group of nanobodies with a high similarity in their CDR3 sequence (identical length and > 80% sequence identity). Nanobodies from the same family derive from the same B-cell lineage and likely bind to the same epitope on the target. Immunizations, library construction, selection by panning and nanobody characterization were performed according to standard procedures (Pardon et al., 2014). Five nanobodies were further characterized.

The nanobodies were expressed in *E. coli* WK6 cells and purified following standard procedures. Specifically, the cell pellet was resuspended in TES buffer (0.2 M TRIS, pH 8, 0.5 mM EDTA, 0.5 M sucrose) supplemented with one protease inhibitor tablet (Roche). Osmotic shock was performed by the addition of diluted TES buffer to release the periplasmic proteins. The solution was first centrifuged for 20 min at 10,000 × g and additionally for 30 min at 100,000 × g. The supernatant was applied to CaptureSelect beads (Thermo Fisher Scientific), which were equilibrated with wash buffer (20 mM NaPi, pH 7.5, 20 mM NaCl). After three column volumes of washing, the nanobody was eluted with 20 mM HEPES, pH 7.5, 1.5 M MgCl_2_. The nanobodies were further purified on a HiLoad 16/600 Superdex 75 pg column in 20 mM HEPES, pH 7.5, 150 mM NaCl, 5 % glycerol, concentrated with a 5 kDa cut-off concentrator, flash-frozen and stored at −80 °C until further use.

### Expression and purification of macrobody 26

The nanobody 26 (Nb26) was first inserted into a pBXNPH3 vector containing a C-terminal penta-histidine tag preceded of a HRV-3C protease recognition sequence. The maltose binding protein (MBP) was then inserted in frame with the 3’ end of the nanobody, with two prolines as a linker between the two genes as described in (Botte et al., 2021). The resulting macrobody (Mb26) was expressed in *E. coli* WK6 cells as above. The cell pellet was resuspended in TES buffer (0.2 M TRIS, pH 8, 0.5 mM EDTA, 0.5 M sucrose) supplemented with one protease inhibitor tablet (Roche). Osmotic shock was performed by the addition of diluted TES buffer to release the periplasmic proteins. The solution was first centrifuged for 20 min at 10,000 × g and additionally for 30 min at 142,000 × g. The supernatant was further purified by immobilized-metal affinity chromatography (IMAC) on a gravity column. The beads were pre-equilibrated in 20 mM NaPi at pH 7.5, 300 mM NaCl, 5% glycerol, 15-30 mM imidazole, 0.5 mM TCEP and incubated. Loaded beads were washed with increasing imidazole concentrations (20 mM NaPi at pH 7.5, 300 mM NaCl, 5% glycerol, 15-30 mM imidazole, 0.5 mM TCEP, 0.03% DDM). The proteins were eluted from the column with a buffer containing high imidazole concentration (20 mM NaPi at pH 7.5, 150 mM NaCl, 5% glycerol, 250 mM imidazole, 0.5 mM TCEP, 0.03% DDM) and combined with 1 mg of 3C protease to perform the His-tag cleavage. The cleaved protein was recovered by negative IMAC, concentrated to 0.5 ml using a 30 kDa concentrator (Corning® Spin-X® UF concentrators) and run on an ÄKTA Pure system (GE Healthcare Life Sciences), using a Superdex 75 Increase 10/300 column. Fractions containing the protein were pooled, concentrated, flash frozen and stored at −80°C until further use.

### Thermal stability measurements

The differential scanning fluorimetry method was used to follow the thermal unfolding event (Kotov et al., 2019) of Nb17, Nb26, Nb38, DtpC, DtpC-Nb17, DtpC-Nb26, DtpC-Nb38, Mb26, DtpC-Mb26, and split sfGFP-DtpC_1-475_-Nb26 with a Prometheus NT.48 device (NanoTemper Technologies, Munich, Germany). The purified proteins were diluted to 16 µM, and the complexes were formed using a 1:1.5 molar ratio of membrane protein: fiducial. The fluorescence at 330 and 350 nm was recorded over a temperature gradient scan from 15° to 95°C and processed in GraphPad Prism 9.0 (GraphPad Software).

### AlphaFold2 predictions

Structures with the following sequences were used as input for AlphaFold2 structure prediction (Jumper et al., 2021), and AMBER relaxation. The best ranked models were used for visualization.

>split sfGFP-DtpC_1-475_

MSKGEELFTGVVPILVELDGDVNGHKFSVRGEGEGDATNGKLTLKFICTTGKLPVPWPTLVTTLTYGVQCFSRYPDHMKRHDFFKSAMPEGYVQERTISFKDDGTYKTRAEVKFEGDTLVNRIELKGIDFKEDGNILGHKLEYNKTPSQPRAIYYIVAIQIWEYFSFYGMRALLILYLTHQLGFDDNHAISLFSAYASLVYVTPILGGWLADRLLGNRTAVIAGALLMTLGHVVLGIDTNSTFSLYLALAIIICGYGLFKSNISCLLGELYDENDHRRDGGFSLLYAAGNIGSIAAPIACGLAAQWYGWHVGFALAGGGMFIGLLIFLSGHRHFQSTRSMDKKALTSVKFALPVWSWLVVMLCLAPVFFTLLLENDWSGYLLAIVCLIAAQIIARMMIKFPEHRRALWQIVLLMFVGTLFWVLAQQGGSTISLFIDRFVNRQAFNIEVPTALFQSVNAIAVMLAGVVLAWLASPESRGNSTLRVWLKFAFGLLLMACGFMLLAFDARHAAADGQASMGVMISGLALMGFAELFIDPVAIAQITRLKMSGVLTGIYMLATGAVANWLAGVVAQQTTESQISGMAIAAYQRFFSQMGEWTLACVAIIVVLAFATRFLFST PNSHNVYITADKQKNGIKANFKIRHNVEDGSVQLADHYQQNTPIGDGPVLLPDNHYLSTQSVLSKDPNEKRDHMVLLEFVTAAGITHGMDELYK

>split sfGFP-DtpC_FL_

MSKGEELFTGVVPILVELDGDVNGHKFSVRGEGEGDATNGKLTLKFICTTGKLPVPWPTLVT TLTYGVQCFSRYPDHMKRHDFFKSAMPEGYVQERTISFKDDGTYKTRAEVKFEGDTLVNRIELKGIDFKEDGNILGHKLEYNKTPSQPRAIYYIVAIQIWEYFSFYGMRALLILYLTHQLGFDDNHAISLFSAYASLVYVTPILGGWLADRLLGNRTAVIAGALLMTLGHVVLGIDTNSTFSLYLALAIIICGYGLFKSNISCLLGELYDENDHRRDGGFSLLYAAGNIGSIAAPIACGLAAQWYGWHVGFALAGGGMFIGLLIFLSGHRHFQSTRSMDKKALTSVKFALPVWSWLVVMLCLAPVFFTLLLENDWSGYLLAIVCLIAAQIIARMMIKFPEHRRALWQIVLLMFVGTLFWVLAQQGGSTISLFIDRFVNRQAFNIEVPTALFQSVNAIAVMLAGVVLAWLASPESRGNSTLRVWLKFAFGLLLMACGFMLLAFDARHAAADGQASMGVMISGLALMGFAELFIDPVAIAQITRLKMSGVLTGIYMLATGAVANWLAGVVAQQTTESQISGMAIAAYQRFFSQMGEWTLACVAIIVVLAFATRFLFSTPTNMIQESNDNSHNVYITADKQKNGIKANFKIRHNVEDGSVQLADHYQQNTPIGDGPVLLPDNHYLSTQSVLSKDPNEKRDHMVLLEFVTAAGITHGMDELYK

>split sfGFP-DtpC_+5Gly_

MSKGEELFTGVVPILVELDGDVNGHKFSVRGEGEGDATNGKLTLKFICTTGKLPVPWPTLVTTLTYGVQCFSRYPDHMKRHDFFKSAMPEGYVQERTISFKDDGTYKTRAEVKFEGDTLVNRIELKGIDFKEDGNILGHKLEYNKTPSQPRAIYYIVAIQIWEYFSFYGMRALLILYLTHQLGFDDNHAISLFSAYASLVYVTPILGGWLADRLLGNRTAVIAGALLMTLGHVVLGIDTNSTFSLYLALAIIICGYGLFKSNISCLLGELYDENDHRRDGGFSLLYAAGNIGSIAAPIACGLAAQWYGWHVGFALAGGGMFIGLLIFLSGHRHFQSTRSMDKKALTSVKFALPVWSWLVVMLCLAPVFFTLLLENDWSGYLLAIVCLIAAQIIARMMIKFPEHRRALWQIVLLMFVGTLFWVLAQQGGSTISLFIDRFVNRQAFNIEVPTALFQSVNAIAVMLAGVVLAWLASPESRGNSTLRVWLKFAFGLLLMACGFMLLAFDARHAAADGQASMGVMISGLALMGFAELFIDPVAIAQITRLKMSGVLTGIYMLATGAVANWLAGVVAQQTTESQISGMAIAAYQRFFSQMGEWTLACVAIIVVLAFATRFLFSTPTNMIQESNDGGGGGNSHNVYITADKQKNGIKANFKIRHNVEDGSVQLADHYQQNTPIGDGPVLLPDNHYLSTQSVLSKDPNEKRDHMVLLEFVTAAGITHGMDELYK

### Cryo-EM sample preparation, data collection, image analysis, and atomic modelling

One hour before vitrification, the purified protein complexes were thawed on ice and run on a Superdex Increase 200 5/150 column in 0.015% DDM, 100 mM NaCl, 10 mM HEPES (pH 7.5), 0.5 mM TCEP in order to remove the excess of empty detergent micelles earlier generated upon sample concentration. The top fraction reached a concentration ranging between 3 and 6 mg/ml, and for each sample, 3.6 μl were applied to glow-discharged gold holey carbon 2/1 300-mesh grids (Quantifoil). Grids were blotted for 4 s at 0 force and 1-s wait time before being vitrified in liquid propane using a Mark IV Vitrobot (Thermo Fisher Scientific). The blotting chamber was maintained at 4°C and 100% humidity during freezing.

All movies were collected using a Titan Krios (Thermo Fisher Scientific) outfitted with a K3 camera and BioQuantum energy filter (Gatan) set to 10 eV. Automated data acquisitions were set using EPU (Thermo Fisher Scientific). The applied defocus ranged between −0.9 µm and −1.8 µm in all datasets.

For DtpC-Nb26 and DtpC-Mb26, movies were collected at a nominal magnification of ×105,000 and a physical pixel size of 0.85 Å, with a 70-μm C2 aperture and 100-μm objective aperture at a dose rate of 19.5 e−/pixel per second. A total dose of 75 e−/Å^2^ was used with 2.8 s exposure time, fractionated in 50 frames. For split sfGFP-DtpC_1-475_-Nb26, movies were collected at a nominal magnification of ×130,000 and a physical pixel size of 0.67 Å, with a 50-μm C2 aperture and 100-μm objective aperture at a dose rate of 19.0 e−/pixel per second. A total dose of 57 e−/Å^2^ was used with 3 s exposure time fractionated in 40 frames.

All movies were motion-corrected using Relion-3.1 (Scheres, 2012; Zivanov et al., 2018) own implementation of MotionCor2 (Zheng et al., 2017). Contrast transfer function parameters were calculated using CTFFIND4 (Rohou and Grigorieff, 2015). and putative particle coordinates were initially defined using CrYOLO (Wagner et al., 2019).

For DtpC-Mb26, 13257 movies were collected, 3,062,337 coordinates were picked and used for 2D averaging and clustering. For split sfGFP-DtpC_1-475_-Nb26, 7602 movies were collected, 1,049,399 coordinates were picked and used for 2D averaging and clustering. For DtpC-Nb26, 24,333 movies were collected, 6,464,070 coordinates were picked and used for 2D averaging and clustering, and 878,428 particles were used in the final 3D reconstruction. Briefly, DtpC-Nb26 dimeric population was clustered using 3D class averaging in Relion3.1 (Scheres, 2012). Particle trajectories and cumulative beam damage were further corrected by Bayesian polishing in Relion3.1 (Zivanov et al., 2019), and the resulting shiny particles were exported to cryoSPARCv3 (Punjani et al., 2017) for further 3D clustering *via* successive heterogeneous refinement cycles using “bad” and “good” volumes as references to denoise the dataset. Non uniform refinement (Punjani et al., 2020), followed by a local refinement using a soft mask around one transporter unit resulted in a 2.7 Å reconstruction of DtpC. The overall resolution was estimated in CryoSPARCv3 using the FSC = 0.143 cutoff. Local resolution estimations were also calculated in CryoSPARCv3 using the 0.5 FSC cutoff. The two half maps were used as inputs to assess various post-processing strategies such as the CryoSPARC’s sharpening tool, DeepEMhancer (Sanchez-Garcia et al., 2020), and Resolve_cryo-em (Terwilliger et al., 2020). The latter led to a slightly better defined contour of the atoms, and was subsequently used for the last atomic-model refinement of DtpC. The initial models of DtpC and Nb26 were generated using AlphaFold2, and refined against the experimental maps; first in Isolde (Croll, 2018), and last in Phenix (Afonine et al., 2018), principally to refine atomic displacement parameters (B-factors) and perform a slight energy minimization while keeping restrains from Isolde’s reference model. Half-maps, and postprocessed maps of the dimeric arrangement and of the focused refinement, as well as the atomic model of DtpC were deposited in the PDB and EMDB as deposition numbers 7ZC2, and EMD-14618. The atomic model of the dimeric DtpC-Nb26 is available upon request.

### Small-angle X-ray scattering data collection and analysis

Synchrotron SAXS data from solutions of DtpC-Nb26 in β-DDM micelles (SEC-SAXS) were collected on the EMBL P12 (Blanchet et al., 2015) beamline at the PETRA III storage ring (Hamburg, Germany), in a buffer consisting of 0.015% DDM, 100 mM NaCl, 10 mM HEPES (pH 7.5), and 0.5 mM TCEP. Sample (10 mg/ml) was injected onto a Superdex Increase 200 10/300 column (Cytiva) and run at 0.5 ml/min at 20°C. 3000 successive 1 second frames were collected using a Pilatus 2M detector at a sample-detector distance of 3.1 m and at a wavelength of λ = 0.124 nm (I(s) vs s, where s = 4πsinθ/λ, and 2θ is the scattering angle). The data were normalized to the intensity of the transmitted beam and radially averaged; the scattering of the solvent-blank was subtracted using CHROMIXS (Panjkovich and Svergun, 2018). Cryo-EM volume maps of DtpC-Nb26 were fit to the scattering data across the low-angle range (shape region only) using EM2DAM (Franke et al., 2017) at a density threshold of 0.1.

## Data visualization

Graphs were generated using GraphPad Prism 9.0 (GraphPad Software). Molecular graphics and analyses performed with UCSF ChimeraX-1.2.5 (Pettersen et al., 2021). Figures were prepared in Adobe Illustrator 2021.

## 6 Author Contributions

Conceptualization: MK, CL

Methodology: MK, GF, HDTM, EP

Investigation: MK, GF, HDTM, EP

Visualization: MK, HDTM

Funding acquisition: DIS, JS, CL

Project administration: CL

Supervision: DIS, JS, CL

Writing—original draft: MK, CL

Writing—review & editing: MK, GF, HDTM, DIS, EP, JS, CL

## 8 Acknowledgements

We thank the Sample Preparation and Characterization facility of EMBL Hamburg for support in this project and the beamlines P13 and P14 at EMBL Hamburg for regular access. We acknowledge Instruct-ERIC and the FWO for their support to the Nb discovery and Saif Saifuzzaman for the technical assistance during Nb discovery. All past and current group members are acknowledged for their input to this manuscript and their efforts to crystallize DtpC over the years.

## 9 Data Availability statement

Half-maps, and post processed maps of the dimeric arrangement and of the focused refinement, as well as the atomic model of DtpC were deposited in the PDB and EMDB as deposition numbers 7ZC2, and EMD-14618.

