## Supplementary information for "Cryo-EM structure of an atypical proton-coupled peptide transporter: Di- and tripeptide permease C"

### *Supplementary Material*

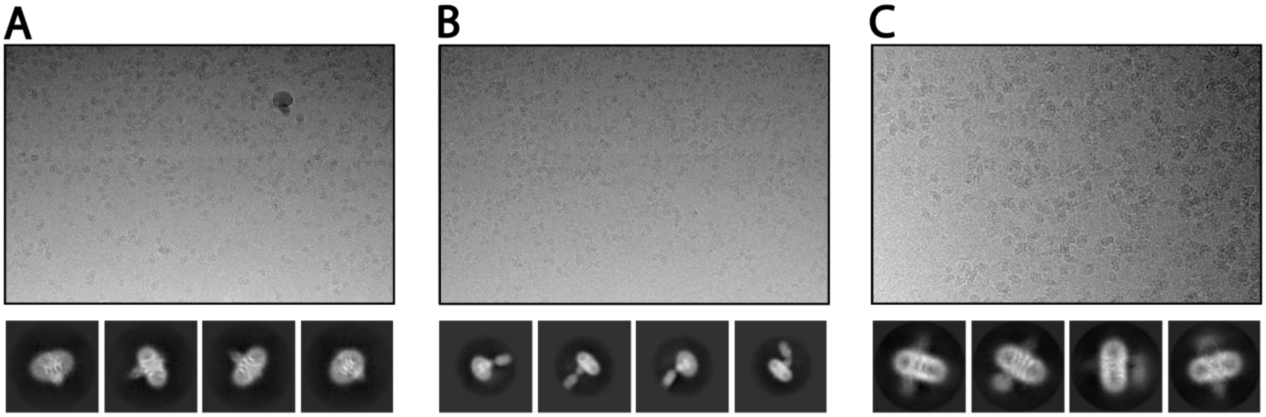


**Supplementary figure 1:** Enlarged representative micrographs and 2D class averages of (A) DtpC-Nb26, (B) DtpC-Mb26, (C) split sfGFP-DtpC1-475-Nb26.

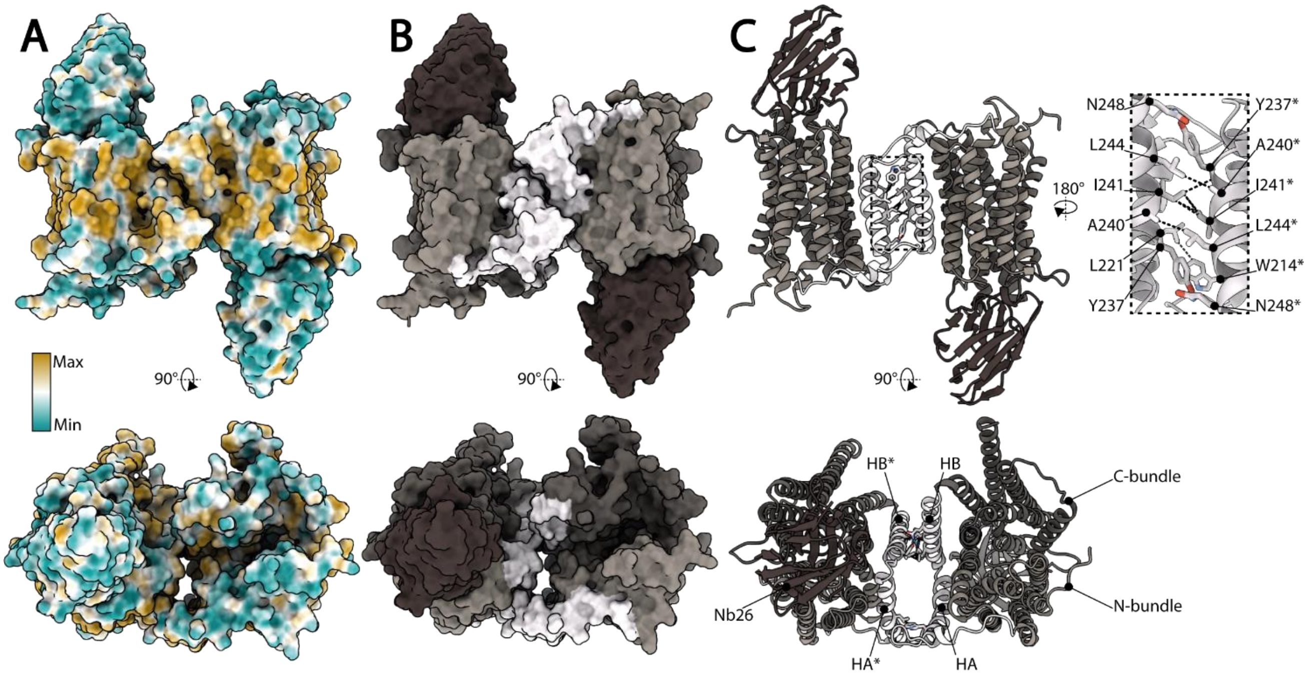


**Supplementary figure 2:** Atomic model of the DtpC-Nb26 inverted dimer represented as surface colored by (A) molecular lipophilicity potential (MLP) and (B) by structural elements labelled in (C), on the ribbon diagram. The close up view on the right shows contacts at the dimer interface between atoms within a 3.8 Å distance.

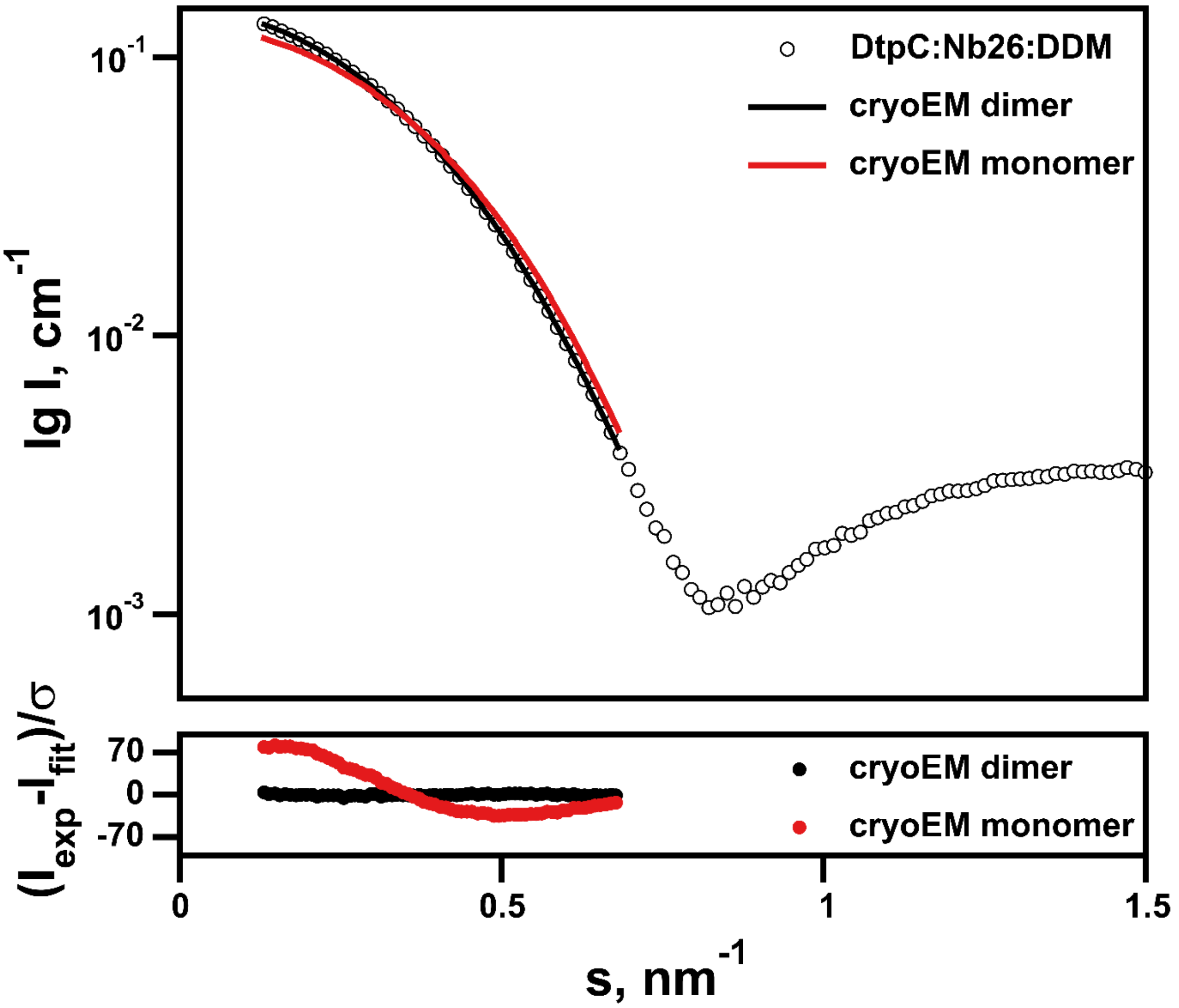


**Supplementary figure 3:** SAXS analysis of cryoEM volume maps of DtpC-Nb26 in the detergent DDM. Upper panel shows the fit of the dimeric (black line) and monomeric (red line) volume maps at a density threshold of 0.1 to the experimental SAXS data (black circles). The residuals of the fits are shown in the lower panel, demonstrating the good fit of the dimeric map and the very poor fit of the monomeric map to the solution SAXS data.
